# Early-life nutrition supplementation and epigenetic age in middle-adulthood among Guatemalan adults

**DOI:** 10.1101/2025.09.25.678652

**Authors:** Melissa Chapnick, Elaine A. Yu, Alicia K. Smith, Karen N. Conneely, Manuel Ramírez-Zea, Zhaohui Qin, Lisa R. Staimez, Viola Vaccarino, Aryeh D. Stein

## Abstract

**Objectives:** Epigenetic clocks are biomarkers of aging. Epigenetic clocks are associated with early-life famine exposure. We investigated the impact of a cluster-randomized early-life nutrition intervention on epigenetic age.

**Methods:** We analyzed follow-up data from participants in the INCAP Nutrition Supplementation Trial, conducted in 4 villages in eastern Guatemala. DNA methylation was measured in buffy coat samples using the Illumina Infinium^TM^ MethylationEPICv2.0 array and standard quality control procedures.

Epigenetic age was quantified using DunedinPACE, PhenoAge, and GrimAge. PhenoAge and GrimAge acceleration were calculated as residuals by regressing epigenetic age on chronological age. We used intent-to-treat difference-in-difference modeling to assess the impact of a protein-energy supplement provided during the first 1,000 days of life (conception to age 2y) on epigenetic age in middle adulthood. Covariates included sex, birth year, the trial supplement type (atole [intervention] vs. fresco [control]), exposure period of supplement (any of the first 1,000 days, other), and a random effect to account for sibships. The primary coefficient of interest was represented by the interaction between supplement type and exposure period.

**Results:** The analysis included 1095 participants (mean age 45.0 y (SD 4.3); 60.3 % female, 40.3 % exposed to any atole during the first 1,000 days, mean DunedinPACE 1.2 (SD 0.1), Phenoage 46.7 y (SD 6.7), and GrimAge 56.3 y (SD 4.1). In difference-in-difference analyses, exposure to atole during any of the first 1,000-day period was associated with lower DunedinPACE (- 0.03, 95% CI -0.06, -0.004), PhenoAge acceleration (- 1.91 y, 95% CI -3.43, -0.39), and GrimAge acceleration (−0.85 y, 95% CI -1.53, -0.11) compared to other exposures. Following additional adjustment for cell type proportions, the direction of the coefficients remained the same but were no longer statistically significant.

**Conclusions:** Exposure to atole during the first 1,000 days was associated with modest reductions in epigenetic age as measured by DunedinPACE, PhenoAge, and GrimAge. These findings complement prior evidence of epigenetic age acceleration among individuals with early-life famine exposure.

## Introduction

In many low- and middle-income countries (LMICs), improvements in public health and nutrition have increased life expectancy [1,2], but this progress is accompanied by a rising burden of age-related diseases [3] that often appear at younger ages and in under-resourced health systems [4]. Early-life nutrition is a key determinant of lifelong health, influencing the risk of accelerated aging and age-related disease [5,6]. Nutrition during sensitive developmental periods—particularly the first 1,000 days from conception to age two—can shape biological systems through processes such as gene expression, metabolic regulation, and immune development [7–9]. These early exposures may influence aging trajectories and heighten vulnerability to age-related diseases [10], challenges that are compounded in LMICs by the “double burden” of malnutrition: the coexistence of early-life undernutrition and adult overnutrition [11–13].

Biological aging is the gradual deterioration of physiological integrity that occurs with time [14]. Biological age differs from chronological age across individuals depending on genetic, environmental, and lifestyle factors [14]. Biomarkers of biological aging allow measurement of these differences and offer insights into factors, such as early-life nutrition, that may influence the aging process [14,15]. Epigenetic clocks provide one such biomarker, by estimating biological age from DNA methylation (DNAm) patterns [14–19]. DNAm-predicted age in excess of chronological age, referred to as epigenetic age acceleration, has been associated with increased risk of chronic diseases, frailty, and earlier mortality [14]. Conversely, younger epigenetic age relative to chronological age - decelerated aging - may indicate resilience to environmental stressors or past exposure to protective early-life environments [14].

Studies of historical famines in the Netherlands[20,21], China[22–25], Ukraine[26], Bangladesh[27], and Nigeria[28,29] have shown that early-life exposure to severe undernutrition is associated with a higher risk of cardiometabolic disease in adulthood. These diseases have, in turn, been linked to accelerated biological aging [30–37]. More recent studies suggest that early-life famine exposure is also associated with faster biological aging [38–41]. Together, this evidence suggests that early-life undernutrition may adversely influence biological aging processes, but to date there is limited evidence that improvements in early-childhood nutrition can reduce the rate of biological aging.

Here we leverage data from the Institute of Nutrition of Central America and Panama (INCAP) Nutrition Supplementation Trial Cohort (INSTC) to test whether improved early-life nutrition influences biological aging, as measured by DNA methylation-based epigenetic clocks. In this cohort, we previously observed that early-life improvements in protein-energy nutrition were associated with a reduced risk of diabetes, an age related disease, in adulthood [42,43]. Whether these nutritional improvements also affect biological aging remains unknown. We hypothesized that individuals exposed to a nutrition supplement during the first 1,000 days of life would have reduced epigenetic age acceleration in adulthood compared to those without such exposure. These analyses aim to assess the biological aging consequences of early-life nutrition and inform strategies to support healthy aging throughout the life course.

## Methods

### Study Design and Population

Participants were enrolled between 1969 and 1977 in a cluster*-*randomized nutrition supplementation trial [44]. Four villages in eastern Guatemala were randomized within pairs to receive either “atole,” a protein-energy supplement or “fresco” a lower-energy, protein-free drink. Atole and fresco had matching micronutrient profiles and were provided ad libitum to village residents. Intake was recorded for pregnant and lactating women and their children (n=2392) up to 7 years of age [44]. Details of the original study design have been previously published [44,45].

Child participants from the original trial have been followed prospectively. We report on data from samples collected during a wave of follow-up conducted in 2015-7 [42,46–52].

### Data Collection

We conducted outreach activities to contact all surviving cohort members residing in Guatemala. All original cohort members were eligible unless they were pregnant or lactating; sessions for these individuals were rescheduled. We collected data between 2015 and 2017 at study clinics located in the original study villages and at INCAP facilities in Guatemala City. Sociodemographic information was obtained through interviews and classified into socioeconomic status tertiles derived from principal component analysis of household characteristics and durable goods, as previously described [42,43]. Fasting venous blood was collected by trained phlebotomists and processed into plasma, serum, and buffy coat aliquots. Immediately after collection, buffy coat aliquots were placed on dry ice. That same day, samples were moved to a −20 °C freezer for temporary storage. They remained there until transport to the INCAP laboratory, where they were transferred to a −80 °C freezer for long-term storage. All samples were then shipped to Emory University, where they remained in -80°C freezers until DNA extraction.

### Ethics Review and Informed Consent

All samples, data collection, and the present analysis, were conducted under protocols approved by the Institutional Review Boards at Emory University (Atlanta, Georgia, USA), and the Institute of Nutrition of Central America and Panama (Guatemala City, Guatemala). Participants provided written informed consent in Spanish prior to data collection.

### DNA Extraction and Methylation Assessment

Genomic DNA was extracted from stored buffy coat samples using the Qiagen QiAmp DNA Mini Kit. We performed initial extractions on the Qiagen QIAcube Automated DNA/RNA Isolation Purification System. We measured DNA concentrations with the Quant-iT^TM^ PicoGreen® dsDNA assay. Samples with low DNA yield were re-extracted manually using the same kit. Extractions took place between August 2023 and March 2024.

To minimize batch and chip effects, our plating strategy balanced 12 exposure groups across each plate, row, and column. We defined groups by exposure type (atole or fresco), exposure timing (full 1,000 days, partial 1,000 days (1 to 999), or none of the first 1,000 days), and biological sex (Male/Female). DNA methylation was quantified at ∼935,000 CpG sites using the Illumina Infinium^TM^ MethylationEPIC v2.0 BeadChip [53,54]. We used the IDOL algorithm [55] to estimate cell type proportions from methylation data using the following workflow: https://github.com/PGC-PTSD-EWAS/EPIC_QC.

Standard pre-processing and quality control for the EPIC v2.0 array followed protocols developed by Pidsley et al. [56]. incorporating techniques from ewastools [57,58] and the Enmix pipeline [59–65]. Quality control procedures aimed to 1) to identify poorly performing samples, 2) to identify poorly performing CpG sites, and 3) to address technical sources of variation in signal. The full workflow is available here: https://github.com/PGC-PTSD-EWAS/EPICv2_pipeline.

### Epigenetic Age Clocks

We calculated epigenetic age using three established DNA methylation clocks: DunedinPACE, PhenoAge and GrimAge [15–17]. DunedinPACE [16] and MethylCIPHER [15,17,66] R packages were used to calculate these clocks. These age clocks were originally developed using data from the Illumina 450k array; however, we used the MethylationEPICv2 array to assay our samples, which does not include all the CpG sites required for clock algorithms [67]. To address this for the DunedinPACE clock, we imputed the mean β-value for each missing CpG site using values from GEO 40279, a publicly available dataset that has been used to train and validate multiple epigenetic clocks [15,18]. For PhenoAge and GrimAge, we applied principal component-based correction methods [68], which reduce technical noise and increase signal to noise ratio in DNA methylation age estimates and are an established method for addressing the differences between the Infinium arrays [67].

We used the PC-Clocks R package to implement principal component corrections [68]. Lastly, we created a variable called *DunedinAge*, which expresses DunedinPACE as an age rather than a rate, allowing direct comparison with PhenoAge and GrimAge. DunedinAge was calculated by multiplying each participant’s DunedinPACE value by their chronological age.

### Exposure to the intervention

Participant exposure to the atole or fresco supplements during the ‘first 1,000 days’ from conception to age two years, was classified based on their birth village, birth date, and an assumed gestation length of 266 days.

Supplementation began on January 1, 1969, in two villages and on May 1, 1969, in the other two villages, ending in all villages on February 28, 1977. We grouped participants into three exposure categories based on their birth date relative to the timing of the trial: (1) full exposure during the first 1,000 days **-** births from September 24, 1969, to February 28, 1975, in two villages, and from January 22, 1970, to February 28, 1975, in two villages; (2) partial exposure (1–999 days)— births from January 1, 1967, to September 23, 1969, in two villages, and to January 21, 1970, in two villages, as well as births after February 28, 1975, in all villages; (3) no exposure during the first 1,000 days – births before January 1, 1967 (Fig 1). We used a difference-in-differences approach to model the interaction between supplement type (atole or fresco) and timing of exposure [42,43].

**Figure 1.**
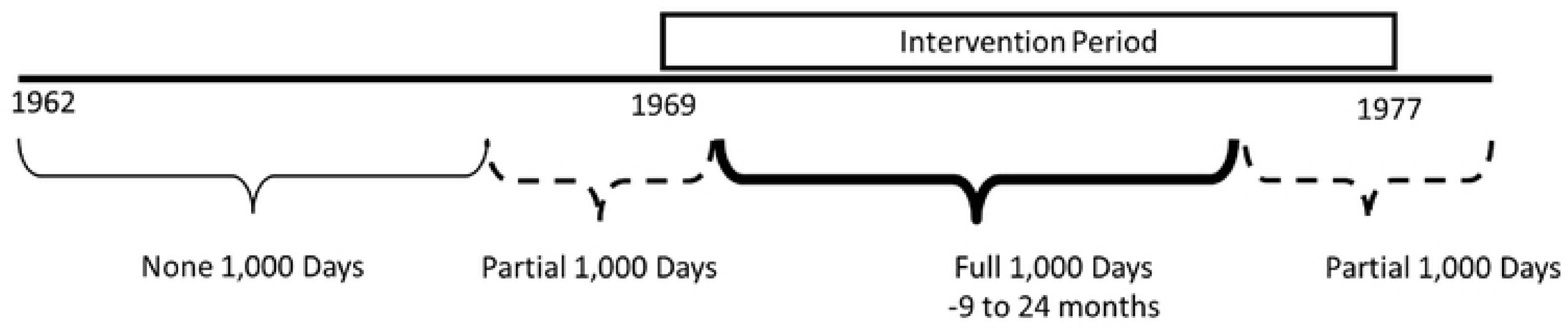
Classification of Nutritional Supplementation Exposure Timing Based on Date of Birth in Relation to the Trial Period.

### Statistical analysis

We used descriptive statistics to characterize the study population and intent-to-treat difference-in-difference modeling to conduct an epigenome-wide association study of the impact of early-life nutritional supplementation on epigenetic age in midlife. We used Pearson correlation to examine relationships among the clocks and between each clock and chronological age.

Because PhenoAge and GrimAge strongly correlate with chronological age, we regressed each on chronological age and used the resulting residuals—which represent the age-independent component of each clock—as the epigenetic age acceleration measures in subsequent analyses. In contrast, DunedinPACE estimates a rate of aging and is not correlated with chronological age; therefore, this clock was analyzed directly, not as a residual. For descriptive purposes, we express DunedinPACE as an age (‘DunedinAge’) by multiplying each participant’s DunedinPACE value by their chronological age.

The primary outcomes of our difference-in-difference models include DunedinPACE, PhenoAge acceleration (residuals), and GrimAge acceleration (residuals). In initial models we categorized exposure into three groups— full 1,000 days, partial 1,000 days, and none—but overlapping confidence intervals between estimates for the full and partial groups led us to combine them into a binary variable (any exposure vs. none) for the final analyses. This binary categorization introduces systematic age differences between participants exposed during the first 1,000 days and those unexposed. To account for this, we created a categorical birth year variable, coding 1962 as 0 and increasing by 1 each year (range 0 – 16), and included this variable and its square to model a quadratic trend [69].

The base model covariates for these analyses include continuous variables for birth year and birth year squared, a binary variable for age of exposure (any first 1,000 days vs. other), a binary variable for treatment assignment (atole vs. fresco), and the interaction term between the binary age of exposure variable and treatment assignment. The primary coefficient of interest is this interaction between any exposure and supplement type compared to other exposure types [42]. Our final model included variables for sex, and a random effect for sibships. These methods are consistent with our previous analyses assessing the impact of early-life nutrition on adult diabetes risk [42]. The final model is as follows:

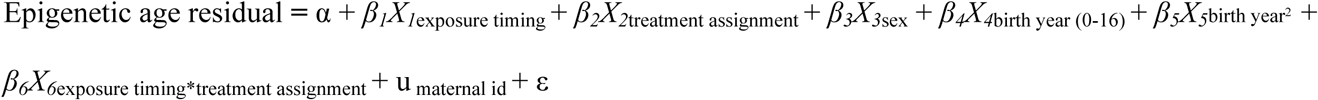

Sensitivity analyses were conducted to assess the robustness of results to adjustments for cell type proportions (CD8T, CD4T, NK, Beta cells, Monocytes), as variation in leukocyte composition can influence DNA methylation profiles and potentially confound associations between exposures and epigenetic outcomes [70]. We also ran sensitivity analyses stratified by sex to assess whether the magnitude and direction of effects differed between men and women.

## Results

A total of 1,095 participants had DNA methylation data available for analysis (Fig 2). The mean age at assessment was 45.0 years (range 37.5y – 55.2y) (Table 1). Approximately 60.3% of the sample were female, consistent with prior waves of follow-up in this cohort. Most participants had completed primary school, and a relatively small proportion had progressed to secondary or higher education. About one-fifth of the participants resided in Guatemala City at the time of follow-up [42,43].

**Figure 2.**
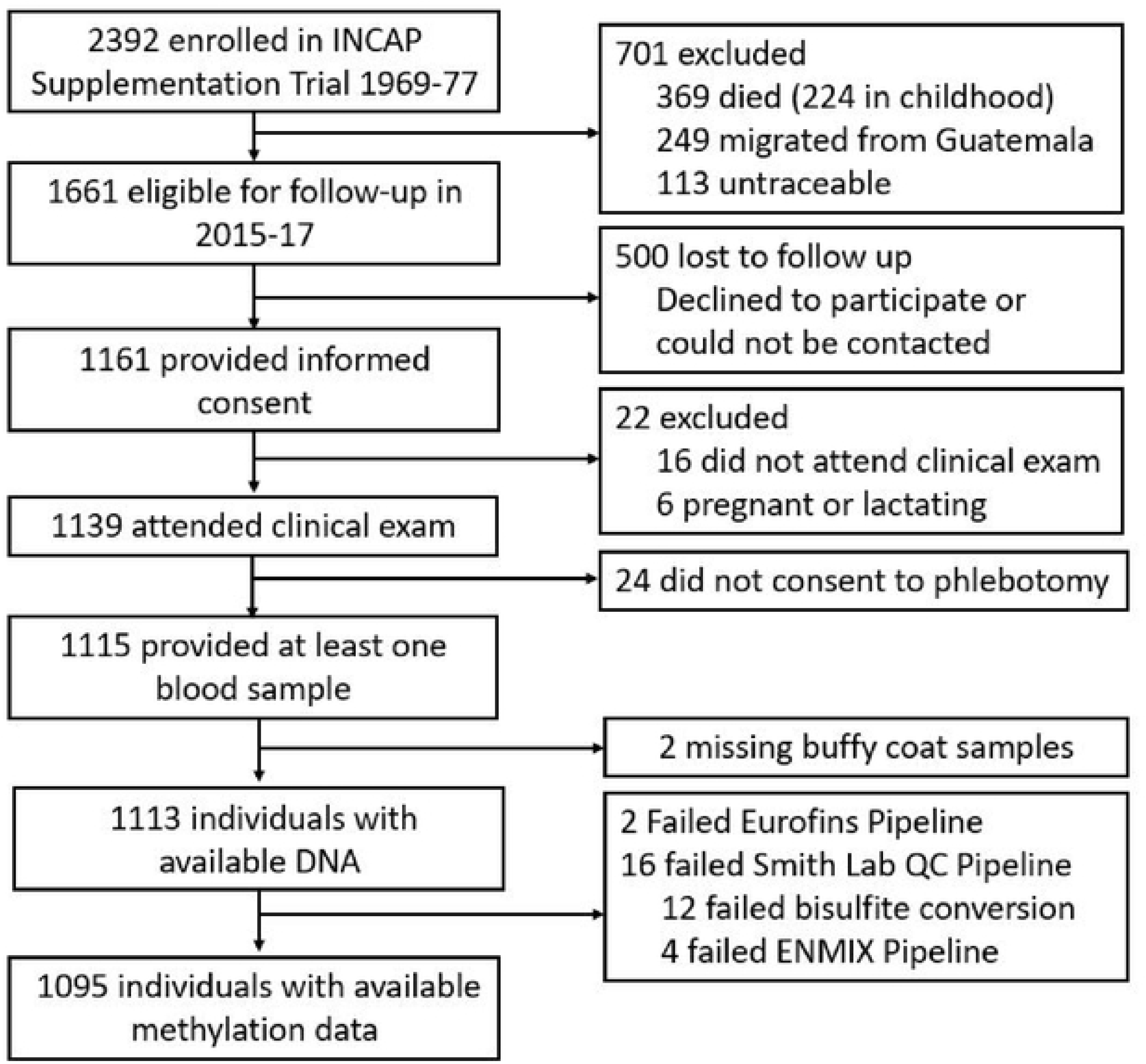
Participant Flow Diagram.

**Table 1.** Selected Characteristics of the Study Population.

| Characteristic | Atole<br>Any 1k Days<br>(n=441) | Other<br>Exposures<br>(n=654) | Pooled<br>(n=1095) |
| --- | --- | --- | --- |
| Age, y | 43.1 (3.1) | 46.3 (4.6) | 45.0 (4.3) |
| Female | 273 (61.9) | 387 (59.2) | 660 (60.3) |
| Socioeconomic Status<sup>2</sup> |  |  |  |
| Poorest | 159 (36.1) | 201 (30.7) | 360 (32.9) |
| Middle | 122 (27.7) | 238 (36.4) | 360 (32.9) |
| Wealthiest | 160 (36.3) | 215 (32.9) | 375 (34.2) |
| Education |  |  |  |
| Less than Primary | 188 (50.5) | 226 (39.0) | 414 (43.5) |
| Primary | 104 (28.0) | 224 (38.7) | 328 (34.5) |
| Some Secondary | 35 (9.4) | 53 (9.2) | 88 (9.3) |
| Secondary and more | 45 (12.1) | 76 (13.1) | 121 (12.7) |
| Residing in Guatemala City | 72 (16.3) | 134 (20.5) | 206 (18.8) |
| Clock Values<sup>3,4</sup> |  |  |  |
| DunedinPACE | 1.2 (0.1) | 1.2 (0.1) | 1.2 (0.1) |
| PhenoAge | 44.8 (6.4) | 48.0 (6.5) | 46.7 (6.7) |
| PhenoAge Residuals<sup>5</sup> | -0.41 (5.94) | 0.28 (5.57) | 0.00 (5.73) |
| GrimAge | 54.7 (3.3) | 57.5 (4.2) | 56.3 (4.1) |
| GrimAge Residuals<sup>5</sup> | -0.26 (2.56) | 0.18 (2.59) | 0.00 (2.59) |

1. Data are mean (SD) or n (%) 2. Socioeconomic status tertiles derived from principal component analysis of household characteristics and durable goods. 3. Based on work from Morgan Levine, the imputation process for CpG sites missing from these clocks inserts the mean value of the CpG site from GSE40279. https://www.ncbi.nlm.nih.gov/geo/query/acc.cgi?acc=GSE40279 After imputation, the clocks were corrected for principal components using the PC-Clocks R package. 4. Epigenetic clocks include DunedinPACE, PhenoAge, and GrimAge, developed as described in Belsky et al. (2022), Levine et al. (2018), and Lu et al. (2019), with principal component corrections to PhenoAge and GrimAge per Higgins-Chen et al. (2022). 5. PhenoAge and GrimAge acceleration were calculated as residuals by regressing epigenetic age on chronological age.

DunedinPACE had a mean value of 1.2, indicating a 20% faster pace of aging compared with the reference value of 1.0 (Table 1, Fig 3). PhenoAge (mean 46.7 y) was 1.7 years higher than the sample’s mean chronological age (45.0 y), whereas GrimAge (mean 56.3 y) was 11.3 years higher (Table 1, Fig 3). The clocks were correlated with chronological age (r = 0.51-0.78), and with each other (r = 0.65 – 0.78) (Fig 4).

**Figure 3.**
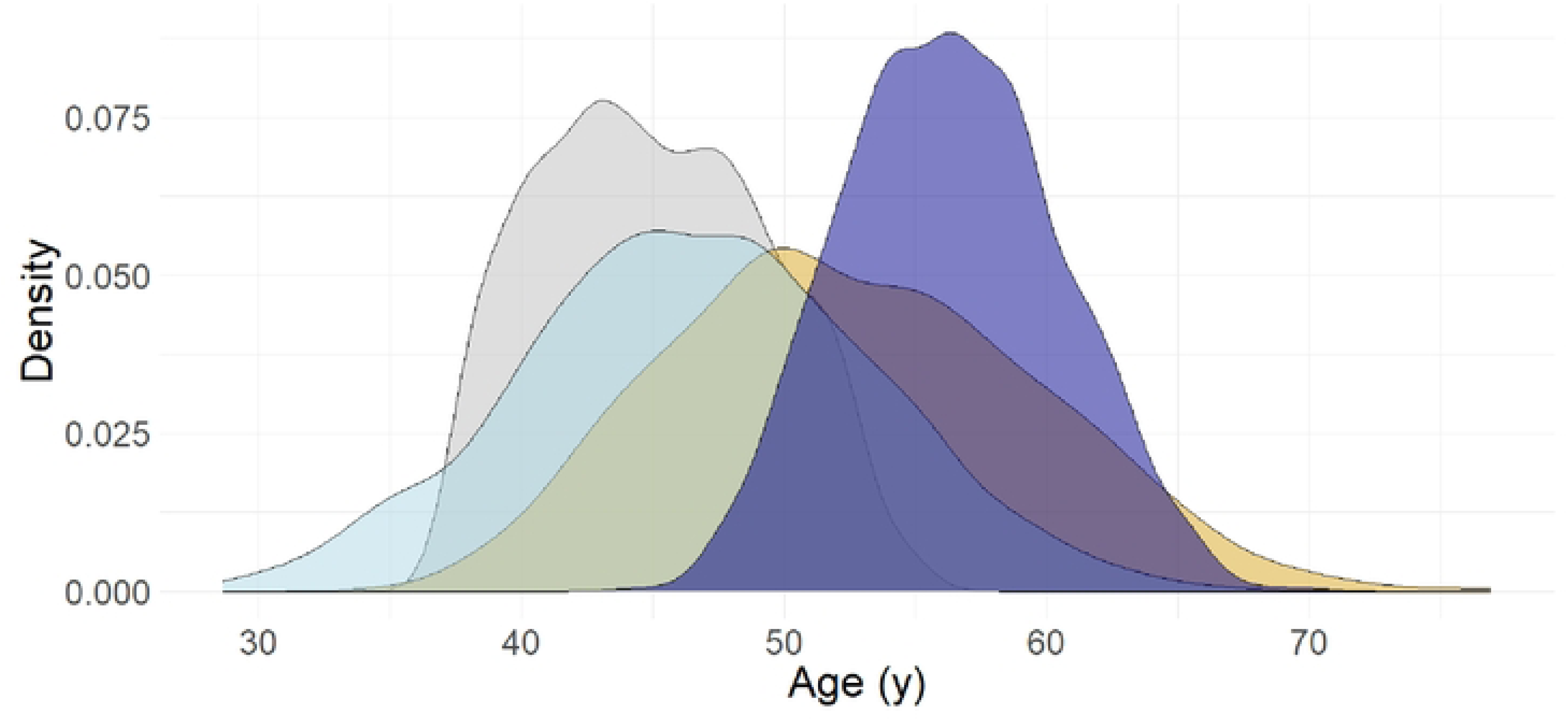
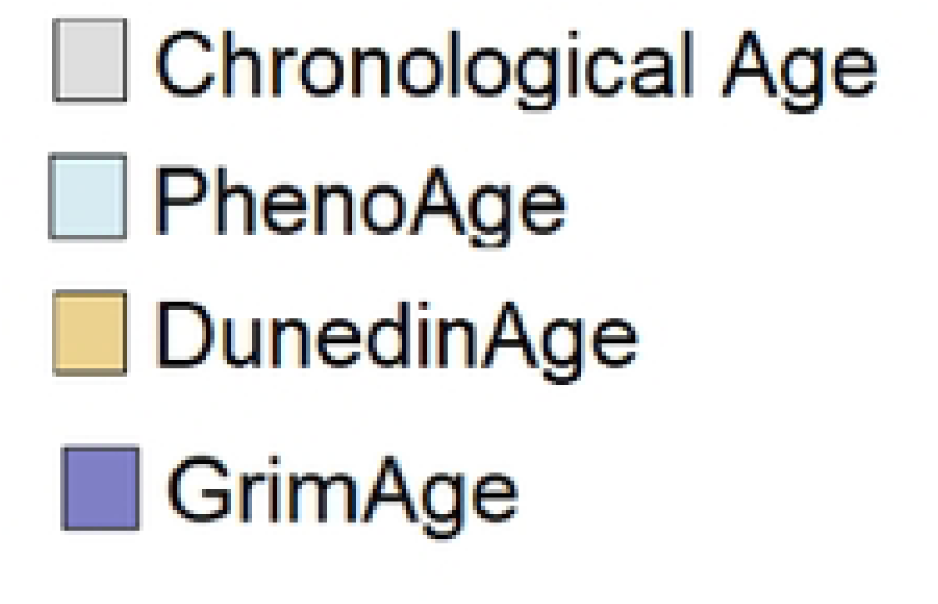
Epigenetic and chronological age distribution of 1095 participants in the INCAP Longitudinal Cohort Study ^1,2^.

**Figure 4.**
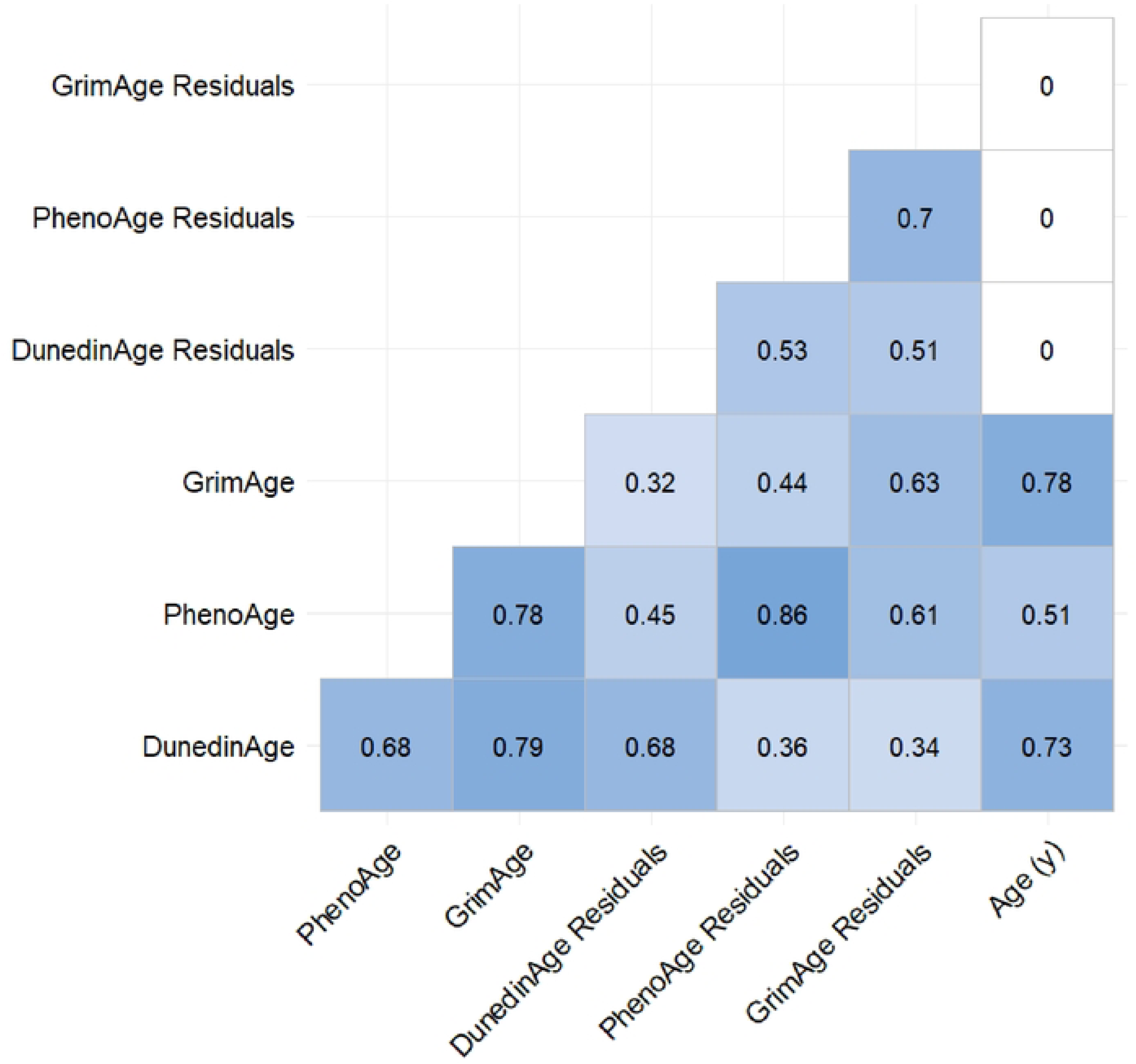
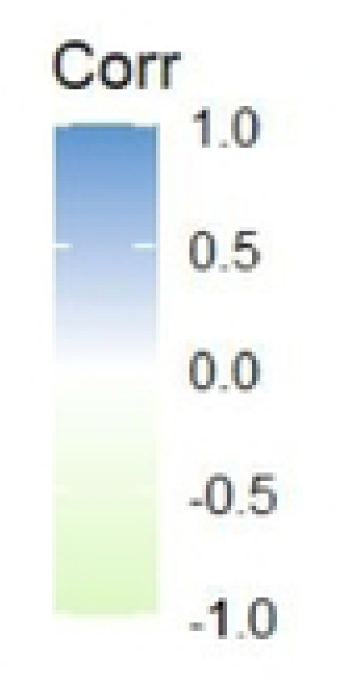
Pearson correlations between chronological age and epigenetic age clocks.

Exposure to atole during any part of the first 1,000 days was significantly associated with lower epigenetic age acceleration across all three clocks (Tables 2 and 3, Fig 5). The effect sizes varied by clock type but were consistent in direction and statistical significance. For example, the DunedinPACE clock suggested that participants exposed to any atole during the first 1,000 days were aging 3% more slowly (∼ 11 days per year) than those with other exposures. Similarly, participants with any atole exposure during the first 1,000 days had PhenoAge residuals that were on average 1.9 years (≈23 months) younger and GrimAge residuals 0.85 years (≈10 months) younger than expected for their chronological age. Measures of association from sensitivity analyses including cell type proportions remained the same in direction but effect sizes were attenuated and no longer statistically significant (Table 3, S1 Fig).

**Figure 5.**
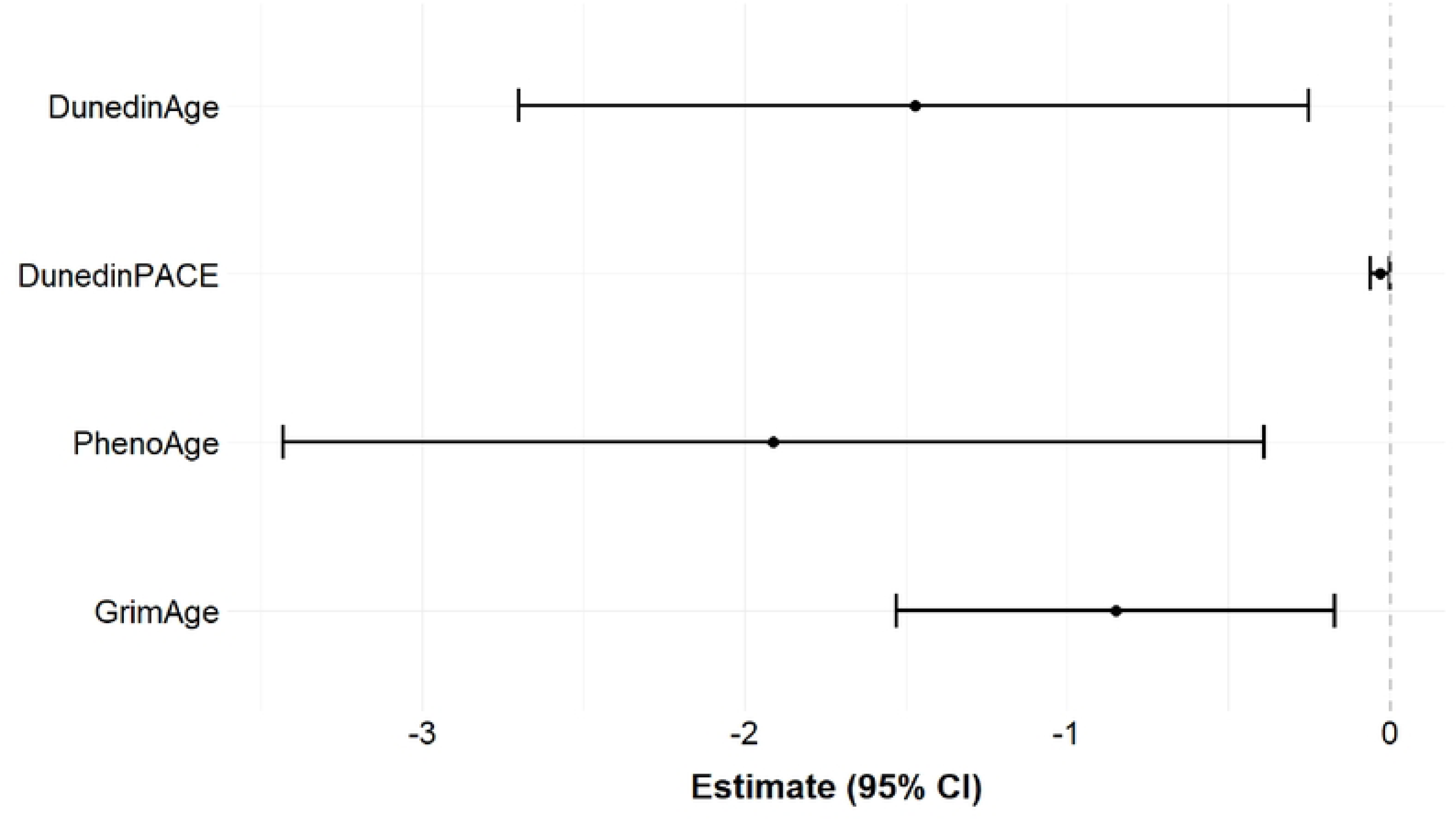
Difference-in-Difference Estimates for Any Atole Exposure During the First 1,000 Days^1,2,3,4^.

**Table 2.**
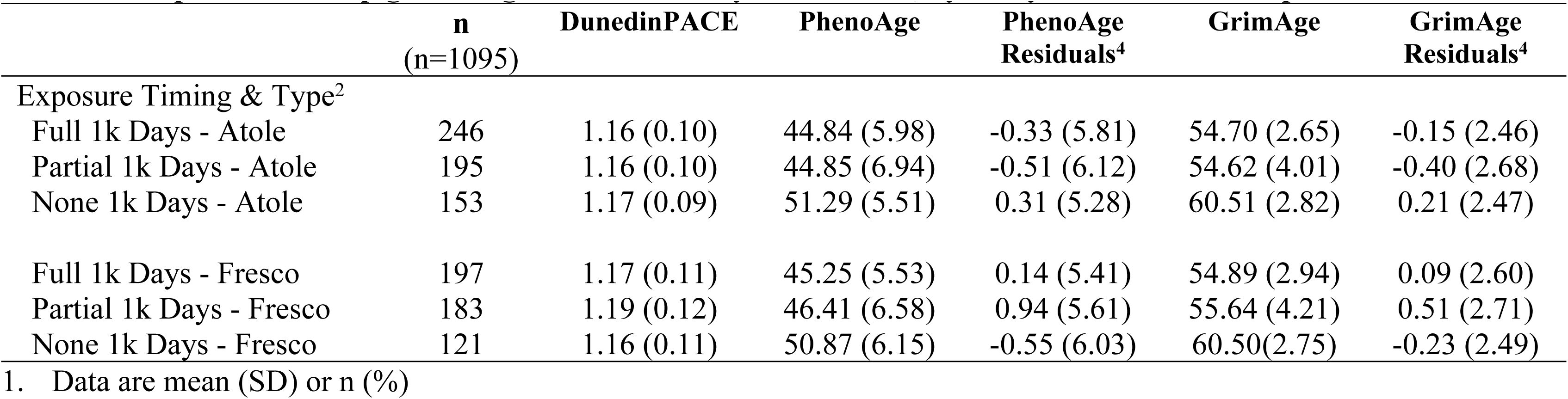
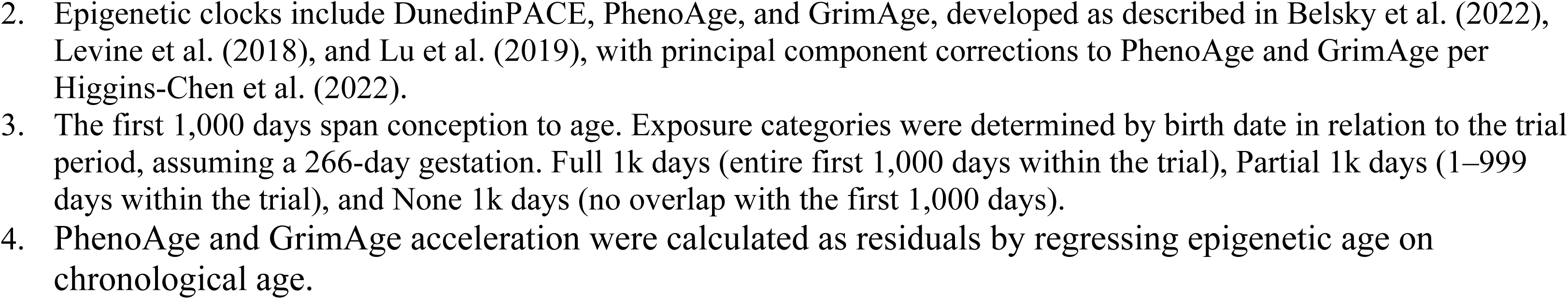
Sample means of epigenetic age as calculated by three clocks, by Early-Life Nutrition Exposures^1,2,3^.

**Figure 5. Difference-in-Difference Estimates for Any Atole Exposure During the First 1,000 Days<sup>1,2,3,4</sup>**
|  | n<br>(n=1095) | DunedinPACE | PhenoAge | PhenoAge<br>Residuals<sup>4</sup> | GrimAge | GrimAge<br>Residuals<sup>4</sup> |
| --- | --- | --- | --- | --- | --- | --- |
| Exposure Timing & Type<sup>2</sup> |  |  |  |  |  |  |
| Full 1k Days - Atole | 246 | 1.16 (0.10) | 44.84 (5.98) | -0.33 (5.81) | 54.70 (2.65) | -0.15 (2.46) |
| Partial 1k Days - Atole | 195 | 1.16 (0.10) | 44.85 (6.94) | -0.51 (6.12) | 54.62 (4.01) | -0.40 (2.68) |
| None 1k Days - Atole | 153 | 1.17 (0.09) | 51.29 (5.51) | 0.31 (5.28) | 60.51 (2.82) | 0.21 (2.47) |
| Full 1k Days - Fresco | 197 | 1.17 (0.11) | 45.25 (5.53) | 0.14 (5.41) | 54.89 (2.94) | 0.09 (2.60) |
| Partial 1k Days - Fresco | 183 | 1.19 (0.12) | 46.41 (6.58) | 0.94 (5.61) | 55.64 (4.21) | 0.51 (2.71) |
| None 1k Days - Fresco | 121 | 1.16 (0.11) | 50.87 (6.15) | -0.55 (6.03) | 60.50(2.75) | -0.23 (2.49) |

1. Data are mean (SD) or n (%)
2. Epigenetic clocks include DunedinPACE, PhenoAge, and GrimAge, developed as described in Belsky et al. (2022), Levine et al. (2018), and Lu et al. (2019), with principal component corrections to PhenoAge and GrimAge per Higgins-Chen et al. (2022). 3. The first 1,000 days span conception to age. Exposure categories were determined by birth date in relation to the trial period, assuming a 266-day gestation. Full 1k days (entire first 1,000 days within the trial), Partial 1k days (1–999 days within the trial), and None 1k days (no overlap with the first 1,000 days). 4. PhenoAge and GrimAge acceleration were calculated as residuals by regressing epigenetic age on chronological age.

**Table 3.** Difference-in-difference estimates comparing ‘Any Atole Exposure’ to ‘other exposures’ During the First 1,000 Days^1,2,3,4^.

|  | n | n<br>sibships |  | DunedinPACE | PhenoAge | GrimAge |
| --- | --- | --- | --- | --- | --- | --- |
| Model 1 Unadjusted | 1095 | -- | □, Any*Type | -0.03 | -1.80 | -0.89 |
|  |  |  | 95% CI | (-0.06, -0.002) | (-3.38, -0.23) | (-1.58, -0.20) |
|  |  |  | p-value | 0.037 | 0.025 | 0.012 |
| Model 2 Fully Adjusted | 1095 | 524 | □, Any*Type | -0.03 | -1.91 | -0.85 |
|  |  |  | 95% CI | (-0.06, -0.00) | (-3.43, -0.39) | (-1.53, -0.17) |
|  |  |  | p-value | 0.023 | 0.014 | 0.015 |
| Model 3 Cell-Type Adjusted | 1095 | 524 | □, Any*Type | -0.02 | -1.02 | -0.50 |
|  |  |  | 95% CI | (-0.04, 0.01) | (-2.08, 0.04) | (-1.01, 0.01) |
|  |  |  | p-value | 0.135 | 0.060 | 0.057 |

1. Epigenetic clocks include DunedinPACE, PhenoAge, and GrimAge, developed as described in Belsky et al. (2022), Levine et al. (2018), and Lu et al. (2019), with principal component corrections to PhenoAge and GrimAge per Higgins-Chen et al. (2022). 2. To remove age-related correlation between epigenetic and chronological age, each clock was regressed on chronological age, and the residuals (age-independent variation) were used as outcomes. 3. Model 1 (unadjusted): any first 1,000-day exposure, supplement type (atole vs. fresco), and an interaction term between any 1,000-day exposure and supplement type. Model 2 (fully adjusted): Model 1’s independent variables and additional adjustment for sex, birth year (0-16), birth year<sup>2</sup>, and a random effect for sibships. Model 3 (cell type adjusted): Model 2’s independent variables and additional adjustment for estimated cell type proportions as a sensitivity analysis. 4. Beta coefficients reflect the interaction between exposure period (any 1,000 days vs. other exposures) and supplement type. For DunedinPACE (null = 1.0), betas indicate the percent change in aging pace. For PhenoAge and GrimAge, beta coefficients represent the difference in epigenetic age acceleration (residuals, in years) associated with atole exposure during the first 1,000 days compared to other exposure period.

Sensitivity analyses considering full and partial exposure separately showed that both full and partial 1,000-day atole exposure were each associated with lower epigenetic age in middle adulthood compared to no exposure during the first 1,000 days. However, only the association for partial 1,000 days was statistically significant, while that for full first 1,000 day exposure was not. The confidence intervals for the estimates for both full and partial groups also overlapped substantially (S1 Table). Sex stratified sensitivity analyses of any 1,000 days vs. none 1,000 days, and full 1,000 days, partial 1,000 days, vs none 1,000 days groups did not reveal any differences in inferences (Tables S2-S5, S1 Fig).

## Discussion

In this study, participants exposed to improved nutrition during the first 1,000 days of life had lower epigenetic age acceleration in middle adulthood compared to those who were not exposed to improved nutrition during the first 1,000 dayss. These results were consistent across three epigenetic age clocks, DunedinPACE, PhenoAge, and GrimAge. Adjusting for cell-type proportions attenuated the associations.

DNAm age estimates and effect sizes varied across clocks, which may reflect the distinct biomarkers used to train each clock [15–17]. PhenoAge was trained to predict multiple aging outcomes, including mortality, cancer, healthspan, physical function, and Alzheimer’s disease [15]. GrimAge was designed to more strongly predict mortality [17], while DunedinPACE measures the pace of aging and was trained on 19 biomarkers spanning cardiovascular, metabolic, renal, hepatic, immune, dental, and pulmonary systems in a birth cohort followed for five decades [16]. When comparing effect sizes across clocks, the patterns are broadly consistent but differ in magnitude. The DunedinPACE results suggest a modest slowing of biological aging (about 3% per year, or roughly 11 days), while PhenoAge and GrimAge residuals indicate larger absolute differences in epigenetic age, with participants exposed to any atole during the first 1,000 days showing an average of nearly 2 years younger (PhenoAge) and just under 1 year younger (GrimAge) than expected. All three clocks showed a consistent direction of association between any atole exposure during the first 1,000 days and epigenetic age. Together, these findings suggest that early-life nutritional supplementation may slow the pace of aging, as captured by DunedinPACE, and reduce cumulative biological aging, as reflected in PhenoAge and GrimAge measures.

### Relationship between early-life nutrition, biological aging, and cardiometabolic disease

In utero famine exposure during the Dutch Hunger Winter demonstrated a similar yet opposing relationship between early-life nutrition and DNAm age in adulthood [41]. DNAm age was accelerated among famine-exposed individuals when measured by DunedinPACE, GrimAge, and PhenoAge, though only DunedinPACE reached significance [41]. In contrast, all three clocks showed significant reductions in DNAm age with exposure to improved early-life nutrition in our analyses. We observed larger effect sizes for PhenoAge and GrimAge than those reported in the famine study. While famine reflects extreme nutritional stress, our findings suggest even modest improvements in nutrition during the first 1,000 days can beneficially influence aging trajectories. In famine studies participants were exposed to severe undernutrition in utero—often receiving fewer than 1,000 kilocalories per day [71,72]. By contrast, participants in the INCAP study received a protein-energy supplement providing approximately 100 additional kilocalories per day throughout early life [44,73]. These modest increases in nutritional intake may result in persistent changes in DNA methylation that regulate genes involved in growth, metabolism, and cellular maintenance, ultimately influencing biological aging decades later [74]. Together, these findings highlight the epigenome’s sensitivity to both severe deprivation and modest improvements in early nutrition.

Early-life nutrition has also been consistently linked to adult cardiometabolic risk, particularly type 2 diabetes (T2DM) [20,25,26,28,42,43]. Famine exposure increases diabetes risk, whereas in this Guatemalan cohort, improved early-life nutrition reduced diabetes risk despite increasing adiposity [42,43]. Cardiometabolic disease itself is associated with accelerated aging, likely through persistent biological mechanisms regulating growth and metabolism [10,74]. This relationship is embedded in second- and third-generation DNA methylation clocks (DunedinPACE, PhenoAge, GrimAge), which strongly predict cardiometabolic outcomes and include CpG sites in inflammatory and metabolic pathways [15–17]. Accelerated epigenetic aging may therefore both reflect cumulative metabolic damage and contribute directly to age-related disease biology [14]. In our cohort, where midlife cardiometabolic disease rates are high (32% obese, 14% diabetic, 38% hypertensive) [42,43], these links are likely complex and bidirectional—epigenetic age may mediate the effects of early nutrition on later health while also being influenced by chronic disease. Understanding how early-life nutrition shapes both biological aging and age-related disease risk is especially important in LMICs, where early undernutrition persists alongside rising chronic disease rates.

Relatedly, sensitivity analyses adjusting for white blood cell composition attenuated the associations between early-life nutrition and epigenetic age acceleration. Immune cell proportions change with chronological age and are influenced by chronic disease [70,75,76]. In populations with high cardiometabolic burden, such as this study sample, cell type proportions may reflect both intrinsic and extrinsic aging processes. Intrinsic aging refers to the natural, well-documented changes in cell type proportions that occur with age, independent of environmental or health-related factors. Extrinsic aging measures (i.e. DunedinPACE, PhenoAge, GrimAge), capture both these natural changes and the cumulative effects of external and health-related influences [14,77]. Because cell proportions themselves are biomarkers of aging, it is expected for extrinsic measures to correlate with them. Therefore, adjusting for cell composition in our analyses may remove meaningful biological variation rather than true confounding, highlighting the complexity of interpreting epigenetic age acceleration in aging populations as it is difficult to disentangle changes due to natural aging from those associated with health- and environment-related aging factors.

### Strengths and Limitations

Our study has several strengths. We leveraged data from a cluster-randomized trial of early-life nutritional supplementation with over five decades of follow-up and high participant retention. The randomized design of the original trial and our use of a difference-in-differences analytical approach strengthens causal inference and minimizes confounding from socioeconomic and household factors and secular trends. We measured DNA methylation using standardized protocols and generated established DNA methylation clocks to estimate epigenetic age.

Several limitations should be noted. Epigenetic age was measured only once, limiting assessment of changes over time and temporality. We previously found that early atole exposure reduced diabetes risk but increased adiposity [42,43]. Since cardiometabolic diseases like diabetes can accelerate biological aging, the single DNAm measurement and high disease burden in this cohort prevent us from determining the direction or causality of these relationships.

Additionally, the epigenetic clocks used in this study were developed primarily in populations of European ancestry whereas our cohort is of Hispanic/Latino origin [15–17]. Although these clocks have shown generalizability across diverse populations, there may still be population-specific biases [78]. We also measured DNAm using the MethylationEPICv2 array, which lacks certain CpG sites used in clocks developed on older arrays. While this may affect comparability, we used established methods to account for the few missing sites.

## Conclusions

In conclusion, early-life exposure to improved nutrition may be associated with lower DNAm age in middle adulthood, supporting the idea that the first 1,000 days are a critical window with lifelong effects on biological aging processes. Early-life nutrition interventions could therefore promote healthy aging and reduce the global burden of age-related disease, particularly in LMICs where early-life undernutrition remains common and populations are rapidly aging. Future research should use repeated measures of DNAm age, diverse populations, and explore mechanistic approaches to clarify how early-life nutrition influences the aging process across the life course.

## Acknowledgements

Dawayland Cobb, Michael Simmond, Seyma Katrinli from Alicia Smith’s lab.

## Notes

**Funding Sources**: This research was supported by NIH R01 Adult epigenetics and telomere length in relation to improved nutrition in early life (R01 DK134509), NIH R01 Early childhood nutrition and adult metabolomic and cardiometabolic profiles (R01 HD075784), and the NIH T32 training grant Multidisciplinary Research Training to Reduce Inequalities in Cardiovascular Health at Emory University (T32 HL130025).

### Competing Interest Statement

The authors have declared no competing interest.

